# Strong inhibitory signaling underlies stable temporal dynamics and working memory in spiking neural networks

**DOI:** 10.1101/2020.02.11.944751

**Authors:** Robert Kim, Terrence J. Sejnowski

**Affiliations:** Computational Neurobiology Laboratory, Salk Institute for Biological Studies, La Jolla, CA 92037, USA; Neurosciences Graduate Program, University of California San Diego, La Jolla, CA 92093, USA; Medical Scientist Training Program, University of California San Diego, La Jolla, CA 92093, USA; Institute for Neural Computation, University of California San Diego, La Jolla, CA 92093, USA; Division of Biological Sciences, University of California San Diego, La Jolla, CA 92093, USA

**Author notes:** Correspondence (R.K.), (T.J.S.).

## Abstract

Cortical neurons process information on multiple timescales, and areas important for working memory (WM) contain neurons capable of integrating information over a long timescale. However, the underlying mechanisms for the emergence of neuronal timescales stable enough to support WM are unclear. By analyzing a spiking recurrent neural network (RNN) model trained on a WM task and activity of single neurons in the primate prefrontal cortex, we show that the temporal properties of our model and the neural data are remarkably similar. Dissecting our RNN model revealed strong inhibitory-to-inhibitory connections underlying a disinhibitory microcircuit as a critical component for long neuronal timescales and WM maintenance. We also found that enhancing inhibitory-to-inhibitory connections led to more stable temporal dynamics and improved task performance. Finally, we show that a network with such microcircuitry can perform other tasks without disrupting its pre-existing timescale architecture, suggesting that strong inhibitory signaling underlies a flexible WM network.

## Introduction

Temporal receptive fields are hierarchically organized across the cortex [1, 2]. Areas important for higher cognitive functions are capable of integrating and processing information in a robust manner and reside at the top of the hierarchy [1–3]. The prefrontal cortex (PFC) is a higher-order cortical region that supports a wide range of complex cognitive processes including working memory (WM), an ability to encode and maintain information over a short period of time [4, 5]. However, the underlying circuit mechanisms that give rise to stable temporal receptive fields strongly associated with WM are not known and experimentally challenging to probe. A better understanding of possible mechanisms could elucidate not only how areal specialization in the cortex emerges but also how local cortical microcircuits carry out WM computations.

Previous experimental studies reported that baseline activities of single neurons in the primate PFC contained unique temporal receptive field structures. Using decay time constants of spike-count autocorrelation functions obtained from neurons at rest, these studies demonstrated that the primate PFC is mainly composed of neurons with large time constants or timescales 1, 6–8. In addition, neurons with longer timescales carried more information during the delay period of a WM task compared to short timescale neurons [8]. Chaudhuri et al. [2] proposed a large-scale computational model where heterogeneous timescales were naturally organized in a hierarchical manner that closely matched the hierarchy observed in the primate neocortex. The framework utilized a gradient of recurrent excitation to establish varying degrees of temporal dynamics [2]. Although their findings suggest that recurrent excitation is correlated with area-specific timescales, it is still unclear if recurrent excitation indeed directly regulates neuronal timescales and WM computations.

Recent experimental studies paint a different picture where diverse inhibitory interneurons form intricate microcircuits in the PFC to execute memory formation and retrieval [9–13]. Both somatostatin (SST) and vasoactive intestinal peptide (VIP) interneurons have been shown to form a microcircuit that can disinhibit excitatory cells via inhibition of parvalbumin (PV) interneurons [14, 15]. Furthermore, SST and VIP neurons at the center of such disinhibitory microcircuitry were causally implicated with impaired associative and working memory via optogenetic manipulations [9, 10, 12, 13]. Consistent with these observations, the primate anterior cingulate cortex, which is at the top of the timescale hierarchy [1], was found to contain more diverse and stronger inhibitory inputs compared to the lateral PFC [16]. A recent theoretical study also showed that inhibitory-to-inhibitory synapses, although much fewer in number compared to excitatory connections, is a critical component for implementing robust maintenance of memory patterns [17].

In order to characterize how strong inhibitory signaling enables WM maintenance and leads to slow temporal dynamics, we constructed a spiking recurrent neural network (RNN) model to perform a WM task and compared the emerging timescales with the timescales derived from the prefrontal cortex of rhesus monkeys trained to perform similar WM tasks. Here, we show that both primate PFC and our RNN model utilize units with long timescales to sustain stimulus information. By analyzing and dissecting the RNN model, we illustrate that inhibitory-to-inhibitory synapses incorporated into a disinhibitory microcircuit tightly control both neuronal timescales and WM task performance. Finally, we show that the primate PFC exhibits signs that it is already equipped with strong inhibitory connectivity even before learning the WM task, implying that a gradient of recurrent inhibition could naturally result in functional specialization in the cortex. We confirm this with our model and show that the task performance of RNNs with short timescales can be enhanced via increased recurrent inhibitory signals. Overall, our work offers timely insight into the role of diverse inhibitory signaling in WM and provides a circuit mechanism that can explain previously observed experimental findings.

## Results

### Spiking recurrent neural network model

To study how stable temporal dynamics associated with WM emerge, we trained a spiking RNN model to perform a WM task. The model used in the present study is composed of leaky integrate-and-fire (LIF) units recurrently connected to one another (see Methods).

The WM task we used to train the spiking RNNs was a delayed match-to-sample (DMS) task (Fig. 1a; see Methods). The task began with a 1 s long fixation period (i.e., no external input) followed by two sequential input stimuli (each stimulus lasting for 0.25 s) separated by a delay period (0.75 s). The input signal was set to either −1 or +1 during the stimulus window. If the two sequential stimuli had the same sign (−1/−1 or +1/+1), the network was trained to produce an output signal approaching +1 after the offset of the second stimulus. If the stimuli had opposite signs (−1/+1 or +1/−1), the network produced an output signal approaching −1.

**Fig. 1.**
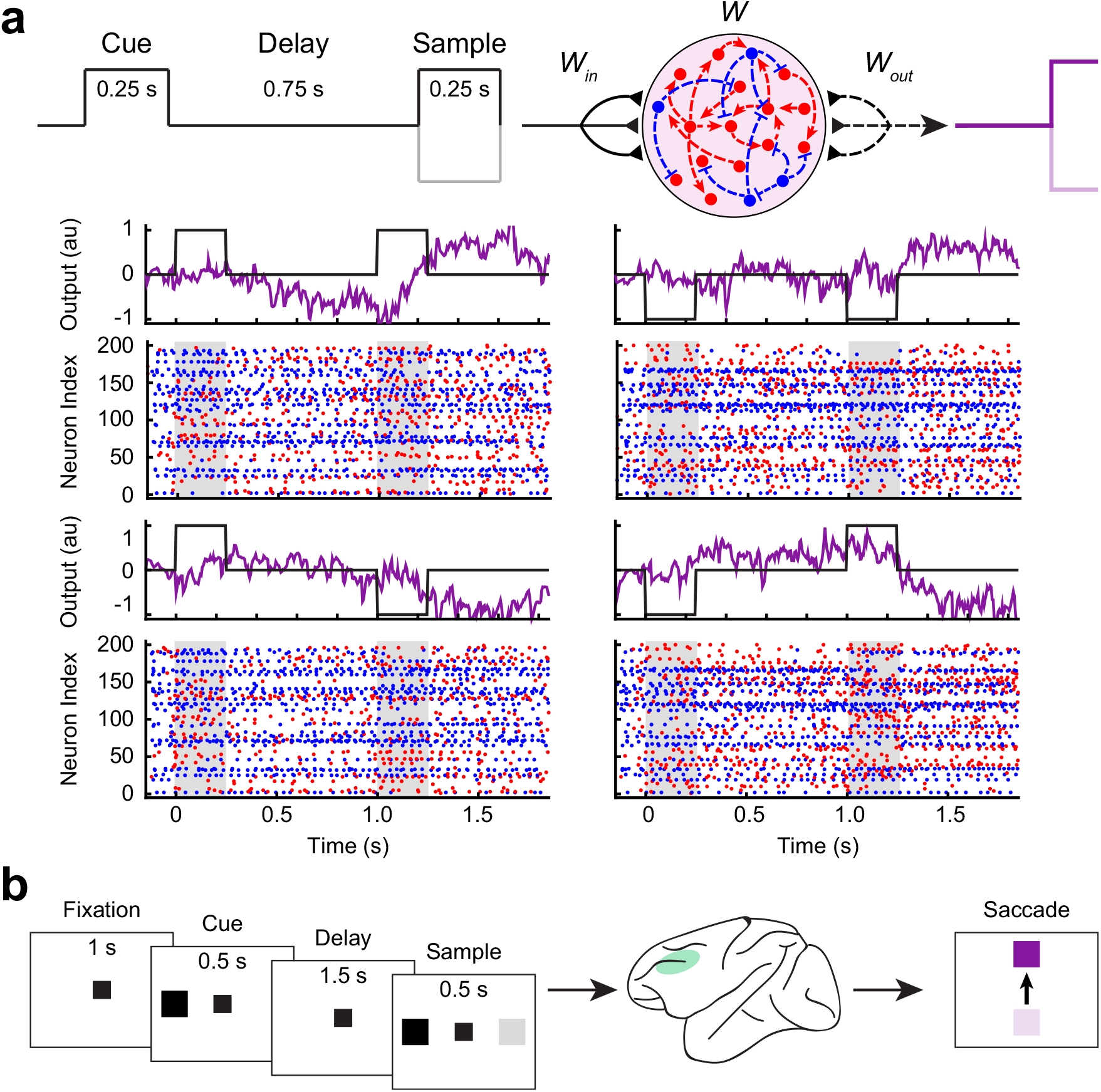
Recurrent neural network model and experimental data. **a**, Spiking recurrent neural network (RNN) model contained excitatory (red circles) and inhibitory (blue circles) units recurrently connected to one another. The model was trained to perform a delayed match-to-sample (DMS) task. Each RNN contained 200 units (80% excitatory and 20% inhibitory), and 40 RNNs were trained to perform the DMS task. The dashed lines (recurrent connections and readout weights) were optimized via a supervised learning method. Example output signals along with the corresponding spike raster plots shown. Gray shading, stimulus window. **b**, Spatial DMS task paradigm used by Constantinidis et al. [18] to train four rhesus monkeys. Extracellular recordings from the dorsolateral prefrontal cortex (green area) were analyzed.

Using a method that we had previously developed, we configured the recurrent connections required for the spiking model to perform the task [19]. Briefly, we trained continuous-variable rate RNNs to perform the task using a gradient descent algorithm, and the trained networks were then mapped to LIF networks. In total, we “trained” 40 LIF RNNs of 200 units (80% excitatory and 20% inhibitory units) to perform the task with high accuracy (accuracy > 95%; see Methods).

### Experimental data

To ensure that our spiking model is a biologically valid one for probing neuronal timescales observed in the cortex, we also analyzed a publicly available dataset containing extracellular spike trains recorded from the dorsolateral prefrontal cortex (dlPFC) of four rhesus monkeys [18, 20, 21]. The monkeys were trained on spatial and feature DMS tasks. A trial for both task types began with a fixation period (1 s in duration) during which the monkeys were required to maintain their gaze at a fixation target. For a spatial DMS trial, the monkeys were trained to report if two sequential stimuli separated by a delay period (1.5 s) matched in spatial location (Fig. 1b). For a feature DMS trial, the monkeys were required to distinguish if two sequential stimuli (in the same spatial location) matched in shape. More details regarding the dataset and the tasks can be found in the Methods and in Qi et al. [20] and Meyer et al. [21].

### Long neuronal timescales in both RNN model and experimental data

Using the spike-count autocorrelation decay time constant as a measure of a neuron’s timescale, previous studies demonstrated that higher cortical areas consist of neurons with long, heterogeneous timescales [1, 7, 8]. Here, we sought to confirm that our spiking RNNs trained on the DMS task and the neural data were also composed of units with predominantly long timescales. For each unit from our RNNs and the dlPFC, we computed the autocorrelation decay time constant (*τ*) of its spike-count during the 1 s fixation period (see Methods) [1]. The baseline activities (average firing rates during the fixation period) of the units that satisfied the inclusion criteria were comparable between the dlPFC data and our model (Fig. 2a; see Methods). Both data contained units with slow temporal dynamics (i.e., long *τ* values) and short *τ* units whose autocorrelation function decayed fast (Fig. 2b). Furthermore, the distribution of the timescales was heavily left-skewed for both data (Fig. 2c,d, left and middle panels) underscoring overall slow temporal properties associated with WM. On the other hand, random RNNs (sparse, random Gaussian connectivity weights) were dominated by units with extremely short timescales (Fig. 2c,d, right panels), suggesting that the long *τ* units observed in the trained RNNs were the result of the supervised training.

**Fig. 2.**
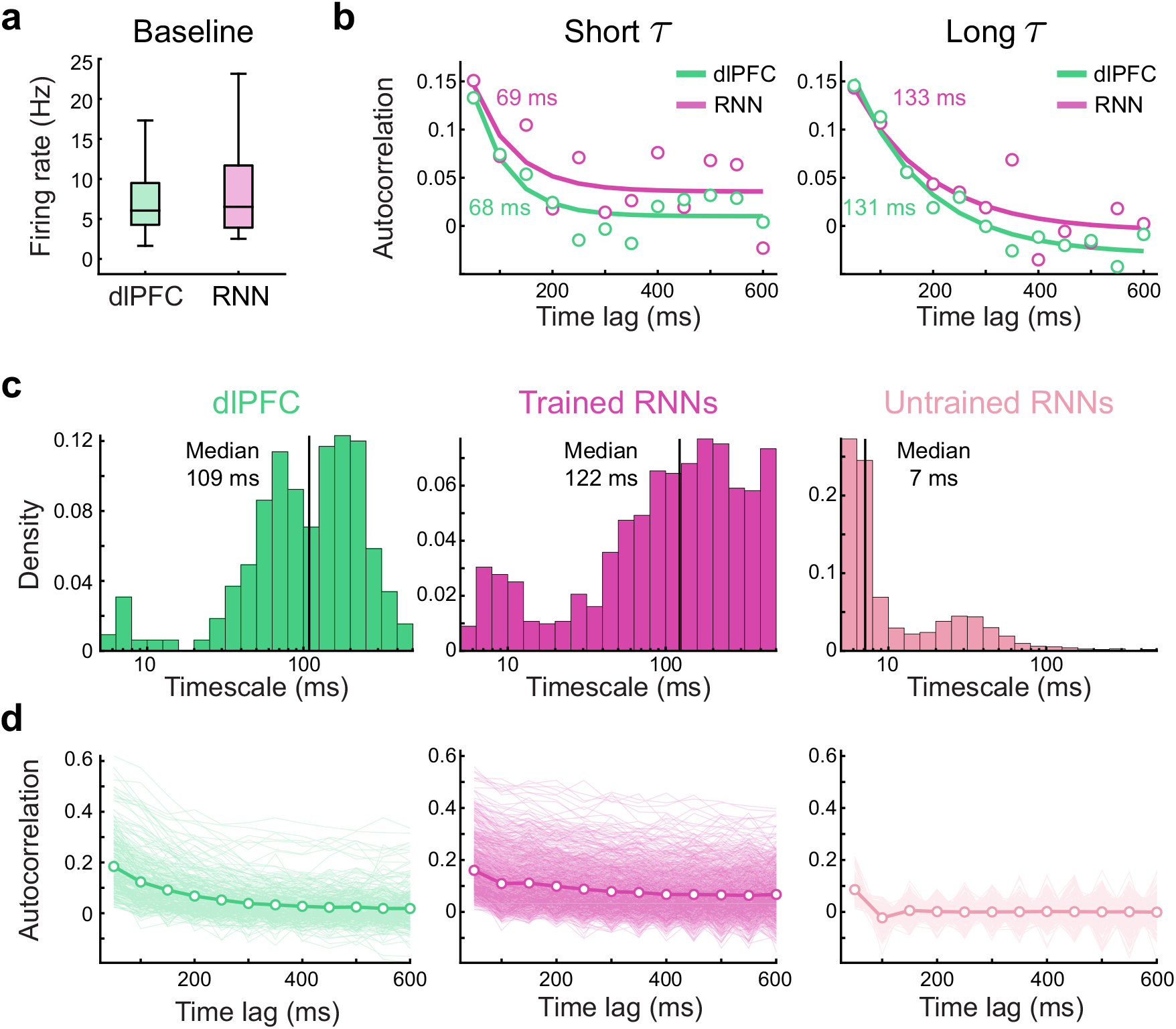
RNN model trained on the DMS task and the dlPFC data contain units with long timescales. **a**, Distribution of the firing rates during the fixation period was not significantly different between the experimental data and the RNN model (*P* < 0.70, two-sided Wilcoxon rank-sum test). **b**, Autocorrelation decay curves from example units with short (left) and long (right) timescale values. **c**, Histograms of the distribution of the timescales from the experimental data (*n* = 325; green), trained RNNs (*n* = 931; magenta), and random RNNs (*n* = 3963; light magenta). Solid vertical lines represent median log(*τ*). **d**, Autocorrelation decay curves from single units (light) and the population average autocorrelation (bold) for the dlPFC data, trained RNNs, and random RNNs. For the random RNNs, only 20% of the total single unit traces shown. Boxplot central lines, median; red circles, mean; bottom and top edges, lower and upper quartiles; whiskers, 1.5*interquartile range; outliers not plotted.

### Long neuronal timescales are essential for stable coding of stimuli

Next, we investigated to see if units with longer *τ* values were involved with more stable coding compared to short *τ* units using cross-temporal decoding analysis [8, 22, 23]. Briefly, for each cue stimulus identity, the trials of each unit were divided into two splits in an interleaved manner (i.e., even vs. odd trials). All possible pairwise differences (in instantaneous firing rates) between cue conditions were computed within each split. Finally, a Fisher-transformed Pearson correlation coefficient was computed between the pairwise differences of the first split at time *t*_1_ and the differences of the second split at time *t*_2_ (see Methods). Therefore, a high Fisher-transformed correlation value (i.e., high discriminability) represents a reliable cue-specific difference present in the network population.

We performed the above analysis on short and long neuronal timescale subgroups from the neural data and the RNN model. A unit was assigned to the short *τ* group if its timescale was smaller than the lower quartile value. The upper quartile was used to identify units with large *τ* values. There were 64 units in each subgroup for the experimental data. For the RNN model, there were 230 units in each subgroup.

The cross-temporal discriminability analysis revealed that stronger cue-specific differences (i.e., higher discriminability) across the delay period were present in the long *τ* subgroup compared to the short *τ* subgroup for both data (Fig. 3a). The significant decodability during the delay period for the dlPFC dataset mainly stemmed from the spatial task dataset (Supplementary Fig. 1). The within-time discriminability (i.e., taking the diagonal values of the cross-temporal decoding matrices) for the long *τ* group was significantly higher than the discriminability observed from the short *τ* group throughout the delay period for the RNN model (Fig. 3b). Although significant within-time discriminability was not observed for the dlPFC data (Fig. 3b, top), Wasmuht et al. [8] reported significant within-time decodability during the delay period in the primate lateral prefrontal cortex, consistent with our model findings.

**Fig. 3.**
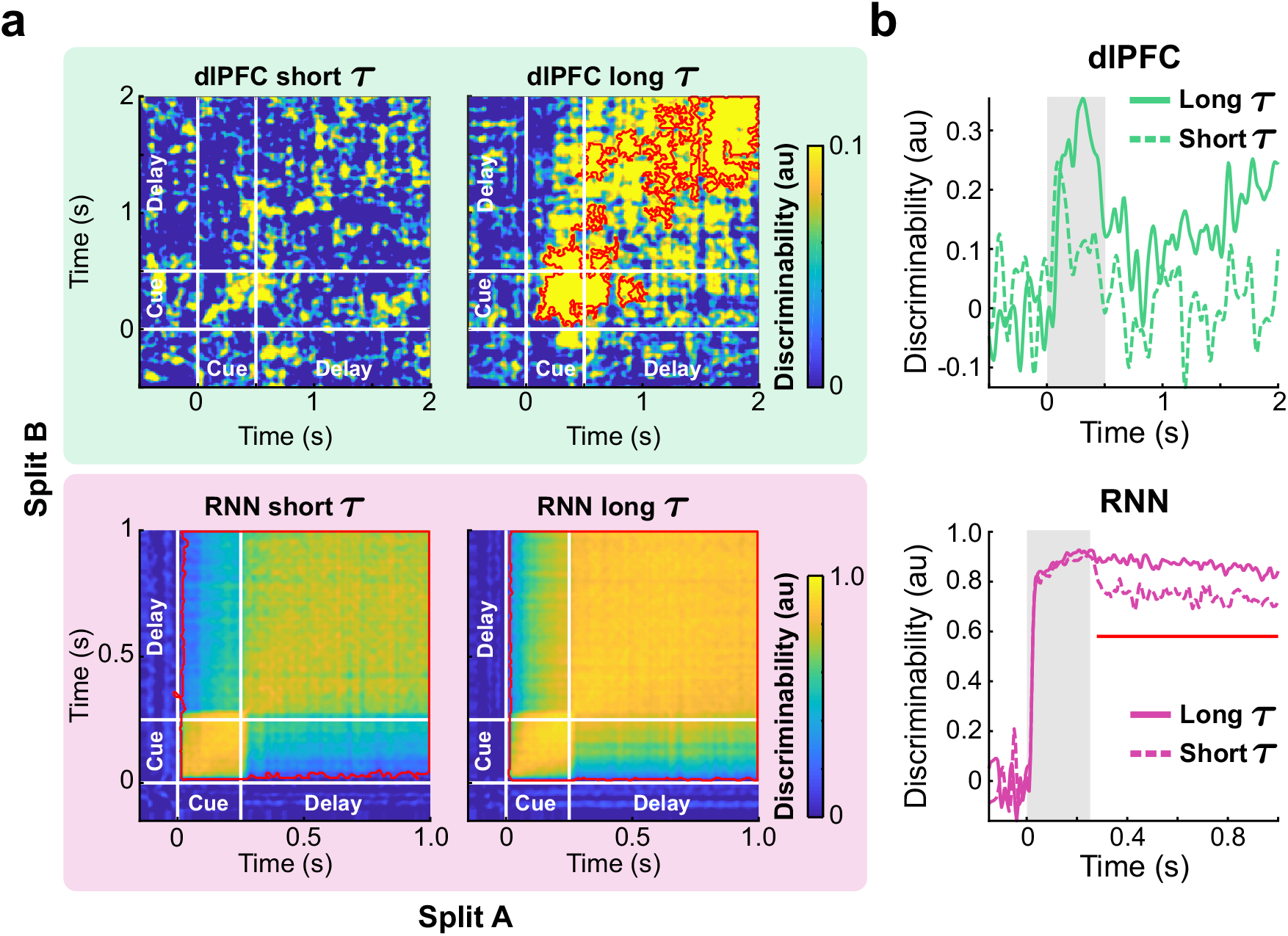
Long *τ* units maintain cue stimulus information during the delay period robustly. **a**, Cross-temporal discriminability matrices for the dlPFC data (top row) and the RNN model (bottom row). Red contours indicate significant decodability (cluster-based permutation test, *P* < 0.05; see Methods). **b**, Within-time discriminability timecourses from the short (dashed) and long (solid) *τ* groups for the dlPFC data and the RNN model. Gray shading, cue stimulus window. Red lines indicate significant differences in decoding between the short and long *τ* groups (cluster-based permutation test, *P* < 0.05; see Methods).

### Strong inhibitory connections give rise to task-specific temporal receptive fields

Neuronal timescales extracted from cortical areas have been shown to closely track the anatomical and functional organization of the primate cortex [1, 2]. For instance, sensory areas important for detecting incoming stimuli house neurons with short timescales. On the other hand, higher cortical areas, including prefrontal areas, may require neurons with stable temporal receptive fields that are capable of encoding and integrating information over a longer timescale.

To investigate if such functional specialization also emerges in our spiking model, we trained another group of spiking RNNs (*n* = 40 RNNs) on a simpler task that did not require WM. The non-WM task, which we refer to as two-alternative forced choice (AFC) task, required the RNNs to respond immediately after the cue stimulus: output approaching −1 for the “−1” cue and +1 for the “+1” cue (Fig. 4a; see Methods). Apart from the task paradigm, all the other model parameters were identical to the parameters used for the DMS RNNs.

**Fig. 4.**
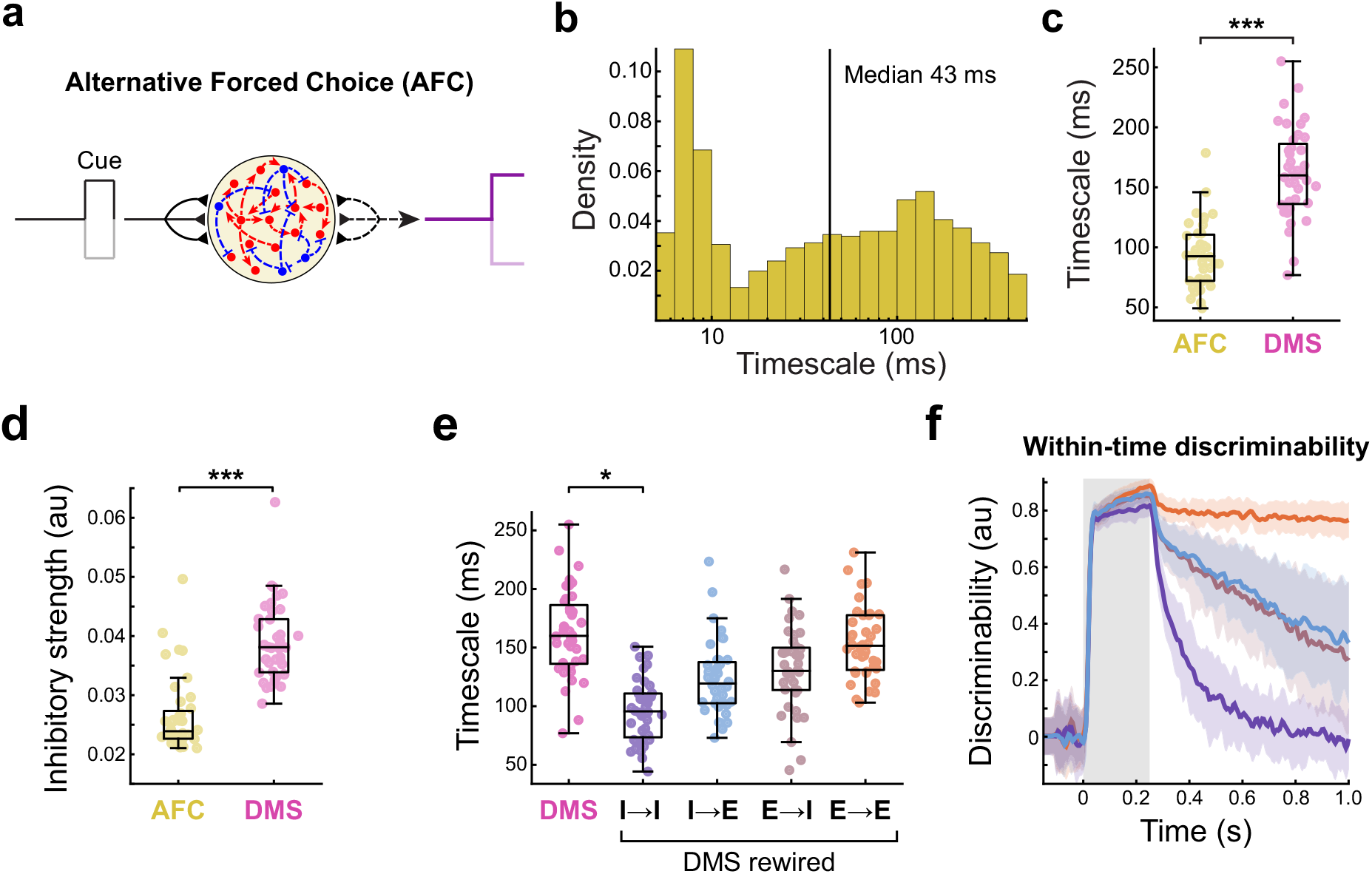
Inhibitory synaptic weights lead to task-specific timescales. **a**, Task paradigm for the alternative forced choice (AFC) task. **b**, Distribution of the neuronal timescales extracted from 40 RNNs trained on the AFC task. Solid vertical line represents median log(*τ*). **c**, Average timescale values from the AFC and DMS RNNs. Each circle represents the average value from one RNN. **d**, Average recurrent inhibitory synaptic strengths from the AFC and DMS models. **e**, Average timescales from the DMS RNNs with each synaptic type rewired randomly (Friedman test, *F* = 74.19, *P* < 0.0001). **f**, Within-time discriminability timecourses averaged across all the DMS RNNs for each rewiring condition. Same color scheme as **e**. The bold line indicates the mean timecourse averaged across 40 RNNs (and all units). Colored shading, ± standard deviation (s.d.). Gray shading, cue stimulus window. Boxplot central lines, median; red circles, mean; bottom and top edges, lower and upper quartiles; whiskers, 1.5*interquartile range; outliers not plotted. \**P* < 0.01, \*\*\**P* < 0.0001 by Wilcoxon signed-rank test (**c**,**d**) or Dunn’s multiple comparisons test (**e**).

Because the AFC task paradigm did not require the RNNs to store information related to the cue stimulus, we expected that these networks would operate on a much faster timescale compared to the DMS RNNs. Consistent with this hypothesis, the AFC RNNs did not contain as many long *τ* units as the DMS RNNs (Fig. 4b), and the timescales averaged by network were also significantly faster for the AFC RNNs (Fig. 4c).

To gain insight into the circuit mechanisms underlying the difference in the timescale distributions of the AFC and DMS RNN models, we compared the recurrent connectivity patterns between these two models. Among other differences, the most notable difference was the inhibitory synaptic strength, which was significantly greater for the DMS RNNs (Fig. 4d). In order to confirm if strong inhibitory signaling led to the long timescales observed in the DMS model, we randomly rewired all the connections belonging to each of the four synaptic types (*I* → *I*, *I* → *E*, *E* → *I*, and *E* → *E*) and computed the timescales again (see Methods). Of the four conditions, only rewiring *I* → *I* synapses resulted in significantly shorter timescales than the timescales from the intact DMS model (Fig. 4e), and the distribution of the timescales pooled from all 40 RNNs with *I* → *I* connections shuffled resembled the distribution obtained from the AFC model (Supplementary Fig. 2). In addition, the amount of cue-specific information maintained during the delay period (as measured by the within-time decoding timecourses) was the lowest for the *I* → *I* rewired condition (Fig. 4f), suggesting that shuffling *I* → *I* synapses was detrimental to memory maintenance.

### Inhibitory-to-inhibitory connections regulate both neuronal timescales and task performance

Given our findings that *I* → *I* connections are important for long neuronal timescales and information encoding, we next investigated if *I* → *I* synapses could be manipulated to provide more stable temporal receptive fields and to improve WM maintenance.

Recent studies revealed that optogenetically stimulating SST or VIP interneurons that specifically inhibit PV interneurons could improve memory retrieval [10–12]. Based on these experimental observations, we expected that strengthening *I* → *I* synapses would increase neuronal timescales and task performance of the DMS RNNs. To test this hypothesis, we first generated another group of RNNs with poor DMS task performance (26 RNNs; mean accuracy ± s.e.m., 71.77 ± 1.43 %). Next, we modeled the effects of optogenetic manipulation of VIP/SST neurons by either decreasing or increasing *I* → *I* synaptic strength (*W*_*I*→*I*_) in each network by 30% (see Methods). Decreasing the connection strength led to significantly shorter timescales compared to the RNNs without any modification (Fig. 5a, left). Strengthening *W*_*I*→*I*_ resulted in a moderate but significant increase in neuronal timescale (Fig. 5a, left). The task performance of the RNNs followed the same pattern: decreasing *W*_*I*→*I*_ severely impaired WM maintenance while increasing *W*_*I*→*I*_ significantly improved task performance (Fig. 5a, right). Increasing *W*_*I*→*I*_ further did not correspond to a significant increase in timescale and task performance (Supplementary Fig. 3). For *I* → *E* connections, only enhancing *W*_*I*→*E*_ resulted in significant changes in both timescale and task performance (Fig. 5b). Manipulating *E* → *I* synapses did not affect the task performance, but decreasing *W*_*E*→*I*_ significantly shortened the timescales (Fig. 5c). Altering the excitatory-to-excitatory connections did not produce any significant changes (Fig. 5d). Overall, these findings suggest that *I* → *I* synapses tightly mediate both temporal stability and WM maintenance. The findings also indicate that the main downstream effect of *I* → *I* connections is to disinhibit excitatory units.

**Fig. 5.**
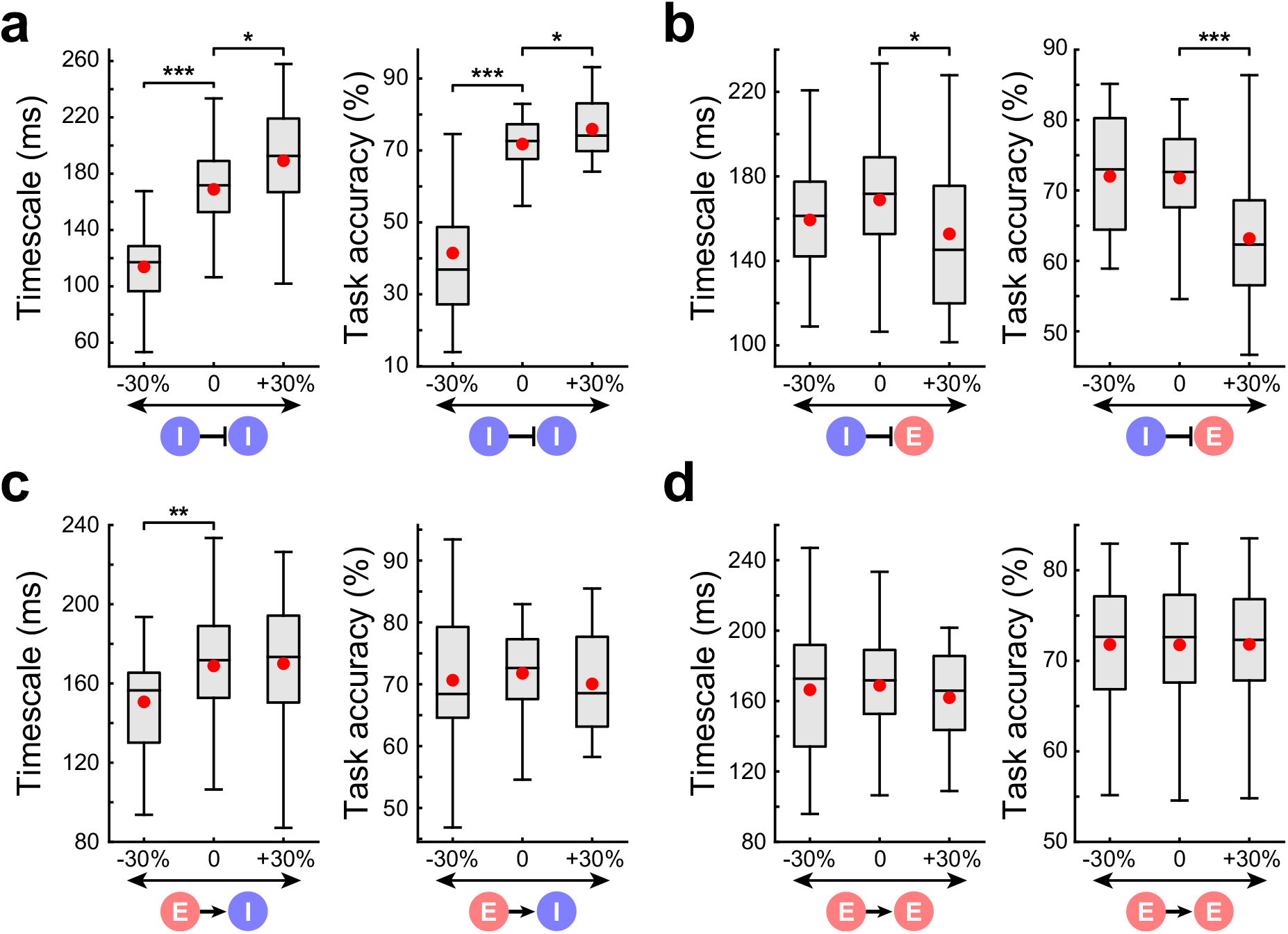
*I* → *I* connectivity strength strongly mediates both neuronal timescales and task performance. **a**, **b**, **c**, **d**, Timescales and task performance changes when *I* → *I* (**a**), *I* → *E* (**b**), *E* → *I* (**c**), or *E* → *E* (**d**) connection strength was either decreased or increased by 30%. Boxplot central lines, median; red circles, mean; bottom and top edges, lower and upper quartiles; whiskers, 1.5*interquartile range; outliers not plotted. \**P* < 0.05, \*\**P* < 0.005, \*\*\**P* < 0.0001 by Wilcoxon signed-rank test.

### Unique inhibitory-to-inhibitory circuitry for WM maintenance

So far, our results indicate that (1) microcircuitry involving specific *I* → *I* connectivity patterns is important for WM (Fig. 4e) and (2) *I* → *I* can be strengthened to enhance both neuronal timescales and task performance (Fig. 5a). Here, we dissect the DMS RNN model to elucidate how specific and strong *I* → *I* connections lead to stable memory retention.

Focusing on inhibitory units only, we first characterized the cue stimulus selectivity from each inhibitory unit in an example DMS network (see Methods). Analyzing the selectivity index values revealed two distinct subgroups of inhibitory units in the network: one group of units favoring the positive cue stimulus and the other group selective for the negative stimulus (Fig. 6a, top). The input weights (*W_in_*) that project to these units closely followed the selectivity pattern (Fig. 6a, bottom).

**Fig. 6.**
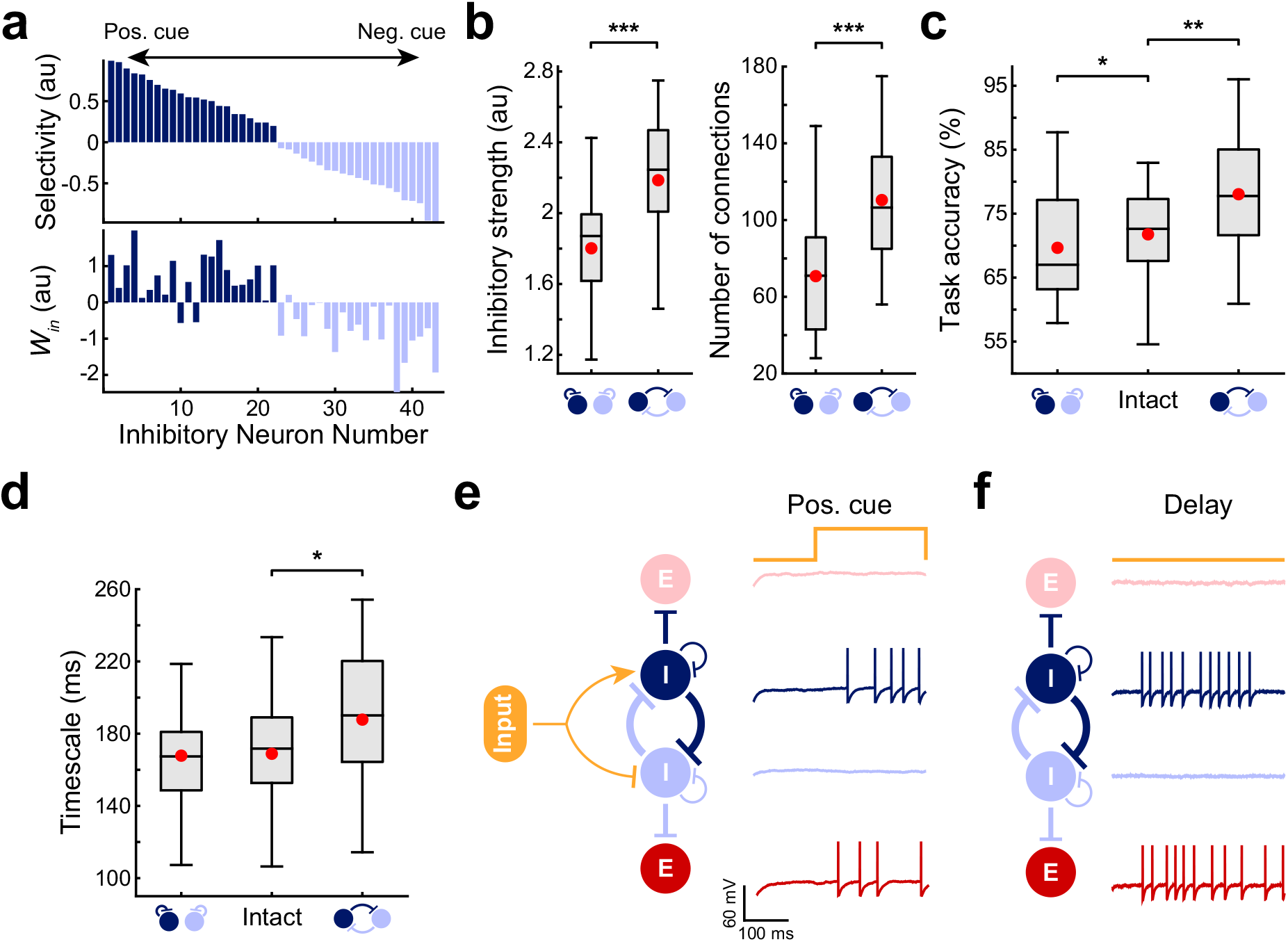
Two oppositely tuned inhibitory subgroups mutually inhibit each other for WM maintenance. **a**, Cue preference selectivity (top) and input weights (*W*_*in*_; bottom) from inhibitory units of an example DMS RNN. The selectivity index values are sorted in descending order. **b**, Average inhibitory strengths (left) and number of inhibitory connections (right) within and across two oppositely tuned inhibitory subgroups from all 40 DMS RNNs. **c**, Average task performance of the DMS RNN model when the within-group or across-group inhibition was increased by 30%. **d**, Average neuronal timescales of the DMS RNNs when the within-group or across-group inhibition was increased by 30%. **e**, **f**, Schematic illustration of the circuit mechanism employed by the DMS RNN model during the cue stimulus window (**e**) and delay period (**f**). The positive cue stimulus was used as an example, and membrane voltage tracings from example units are shown. Dark blue and dark red units indicate units that prefer the positive cue stimulus, while the light blue and light red units favor the negative cue. For simplicity, only recurrent inhibitory connections are shown. Boxplot central lines, median; red circles, mean; bottom and top edges, lower and upper quartiles; whiskers, 1.5*interquartile range; outliers not plotted. \**P* < 0.05, \*\**P* < 0.005, \*\*\**P* < 0.0001 by two-sided Wilcoxon rank-sum test (**b**) or Wilcoxon signed-rank test (**c**, **d**).

Given these two subgroups with distinct selectivity patterns, we next hypothesized that mutual inhibition between these two groups (across-group inhibition) was stronger than within-group inhibition. Indeed, inhibition between the oppositely tuned inhibitory populations was significantly greater (both in synaptic strength and number of connections) than inhibition within each sub-group across all RNNs (Fig. 6b). To confirm that the behavioral improvement we observed with *I* → *I* enhancement in Fig. 5a was largely due to the strengthened across-group inhibition, we increased across-group and within-group *I* → *I* connections separately (see Methods). The DMS RNN performance improved following enhancement of the across-group inhibition, while increasing the within-group inhibition impaired performance (Fig. 6c). In addition, across-group *I* → *I* enhancement resulted in a significant increase in neuronal timescale (Fig. 6d).

In summary, these findings imply that robust inhibition of oppositely tuned inhibitory sub-populations is critical for memory maintenance in our RNN model. For example, a positive cue stimulus activates the inhibitory subgroup selective for that stimulus and deactivates the negative stimulus subgroup (Fig. 6e). Through disinhibition, a group of excitatory units that favor the positive cue stimulus also emerges. During the delay period, the inhibition strength between these two inhibitory subgroups dictates the stability of the cue-specific activity patterns generated during the stimulus window (Fig. 6f).

### Circuit mechanism for WM generates units with long neuronal timescales

The circuit mechanism (Fig. 6e,f) explains why enhancing *I* → *I* connections results in improved WM performance, but it is still not clear how this same mechanism also produces units with long timescales.

Here, we first demonstrate that a high trial-to-trial spike-count variability during the fixation period could give rise to slow decay of the spike-count autocorrelation function. If a neuron exhibits highly variable activity patterns across many trials such that it is highly active (i.e., persistent firing) in some trials and relatively silent in other trials, the Pearson correlation between any two time bins within the fixation window could be large (Fig. 7a). On the other hand, firing activities with a low trial-to-trial variability could result in a weak correlation between two time bins. To directly test this positive relationship between trial-to-trial variability and neuronal timescales, we computed spike-count Fano factors (spike-count variance divided by spike-count mean across trials; see Methods) for the short and long *τ* subgroups in both neural and model data. The Fano factor values for the short timescale subgroup were significantly smaller than the values obtained from the long *τ* group for both data (Fig. 7b). There was also a significant positive correlation between the spike-count Fano factors and neuronal timescales across all the units in both data (Spearman rank correlation, *r* = 0.25, *P* < 0.0001 for dlPFC; *r* = 0.28, *P* < 0.0001 for RNN; Supplementary Fig. 4).

**Fig. 7.**
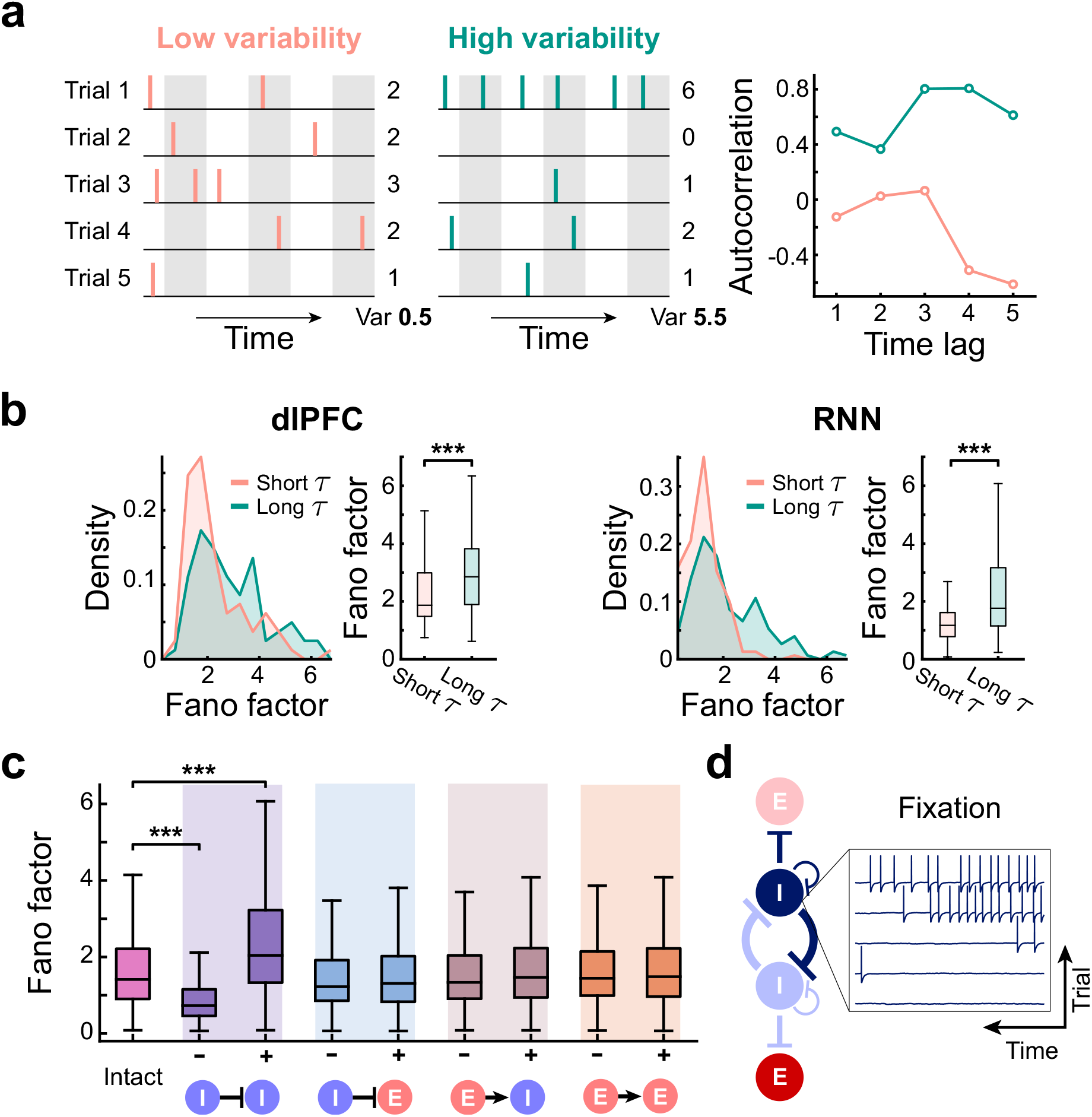
High trial-to-trial spike-count variability during the fixation corresponds to long neuronal timescale. **a**, Schematic illustrating how high spike-count variability across multiple trials can result in a slow decay of the autocorrelation function. **b**, Comparison of the spike-count Fano factors from the short and long *τ* groups in the neural data (left) and the DMS RNN model (right). **c**, Average Fano factors from the DMS model with each of the synaptic type either decreased (“−”) or increased (“+”) by 30% (Kruskal-Wallis test, *H* = 665.2, *P* < 0.0001). **d**, Spiking activity of an example inhibitory unit during the fixation period across 5 trials. The trials were sorted by the number of spikes. Units that were strongly modulated by the disinhibitory circuit mechanism showed highly dynamic baseline firing patterns across trials. Boxplot central lines, median; red circles, mean; bottom and top edges, lower and upper quartiles; whiskers, 1.5*interquartile range; outliers not plotted. \*\*\**P* < 0.0001 by two-sided Wilcoxon rank-sum test (**b**) or Dunn’s multiple comparisons test (**c**).

Manipulating each of the four synaptic types (decreasing or increasing synaptic strength by 30%) in our DMS RNN model revealed that *I* → *I* connections strongly modulated the spike-count Fano factors (Fig. 7c). Enhancing *I* → *I* synaptic strength led to units with more variable spiking patterns across trials, whereas reducing the strength resulted in smaller Fano factors (example shown in Supplementary Fig. 5).

In our RNN model, strong *I* → *I* synapses could give rise to both excitatory and inhibitory units behaving in a highly variable manner during the fixation period (Fig. 7d). For instance, an inhibitory unit selective for the positive stimulus could be partially activated in some trials by chance (i.e., via random noise during the fixation period), and this, in turn, could silence a portion of the negative stimulus inhibitory population (light blue circle in Fig. 7d). This leads to variable firing activities across trials in inhibitory units. Furthermore, the dynamic activity of the inhibitory population could be transferred to the excitatory population via disinhibition. Therefore, *I* → *I* connections play a central role in conferring the network with highly dynamic baseline firing patterns, which then translate to high *τ* values.

### Strong *I* → *I* is an intrinsic property of prefrontal cortex

Finally, we wanted to investigate whether microcircuitry with strong inhibitory-to-inhibitory synapses emerges via learning-related plastic changes. Because extensive plastic changes could disrupt neurons with stable temporal receptive fields, we reasoned that prefrontal cortical areas and other higher cognitive areas are endowed with strong *I* → *I* connections whose connectivity patterns do not undergo significant plastic changes during learning. Instead, learning-related changes occur to the connections stemming from upstream networks that project to these areas. In order to test this hypothesis, we extracted neuronal timescales from the recordings obtained from the same monkeys before they were trained to perform the DMS tasks. In this passive paradigm (Fig. 8a), the monkeys were trained to maintain their gaze at a central fixation point throughout the trial regardless of the stimuli presented around the fixation point [24].

**Fig. 8.**
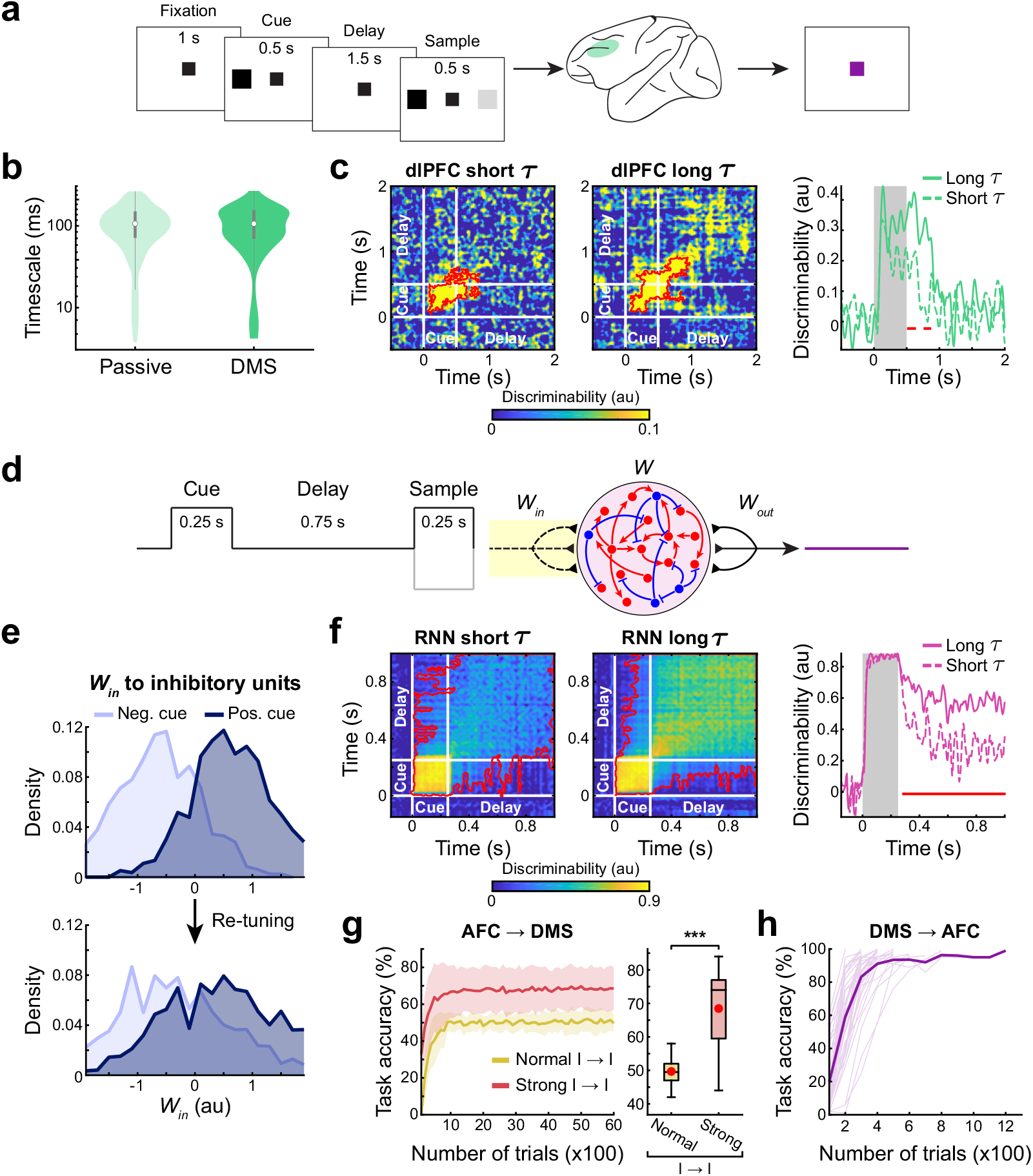
Strong *I → I* connections intrinsic to prefrontal cortex. **a**, Passive task paradigm used by Constantinidis et al. [18] to train the same four monkeys before they learned the DMS tasks (Fig. 1b). **b**, Distribution of the neuronal timescales from the monkeys before (i.e., passive) and after they learned the DMS tasks. **c**, Cross-temporal decoding matrices and within-time decoding timecourses from the short and long *τ* subgroups. **d**, Passive task paradigm used to re-train our DMS RNNs. Only the input weights (dashed lines with yellow shading) were trained. **e**, Distribution of the input weights projecting to the two inhibitory subgroups tuned to the two cue stimuli from all 40 DMS RNNs before (top) and after (bottom) retraining. **f**, Cross-temporal decoding matrices and within-time decoding timecourses from the short and long *τ* subgroups for the re-trained DMS RNNs. **g**, Task performance during re-training of the AFC rate RNNs to perform the DMS task (left) and average performance at the end of training (right). The task performance significantly increased when *I* → *I* connections were strengthened (orange; see Methods). Shaded area, **h**, Task performance during re-training of the DMS rate RNNs to perform the AFC task. Individual networks (light) and mean across 40 DMS RNNs (bold). Boxplot central lines, median; red circles, mean; bottom and top edges, lower and upper quartiles; whiskers, 1.5*interquartile range; outliers not plotted. Red contours indicate significant discriminability (cluster-based permutation test, *P* < 0.05; see Methods). Red lines indicate significant differences in decoding between the short and long *τ* groups (cluster-based permutation test, *P* < 0.05; see Methods). \*\*\**P* < 0.0001 by Wilcoxon signed-rank test.

Surprisingly, the timescales from the spike-train data from the dlPFC of the same four monkeys that learned the passive task were similar to the timescales obtained after the monkeys learned the DMS task (Fig. 8b). In addition, the cue-specific information maintenance during the delay period by long *τ* units was largely abolished, and the within-time decoding was similar between long *τ* and short *τ* neurons (Fig. 8c). These findings suggest that the primate dlPFC was already equipped with stable temporal receptive fields and that learning the DMS task resulted in long *τ* neurons carrying more information during the delay period while preserving the network temporal dynamic architecture.

Next, we asked if we could only optimize the upstream connections (i.e., input weights; *W_in_*) of the DMS RNNs to perform a passive version of the DMS task (Fig. 8d; see Methods). By freezing the recurrent connections (*W*), we ensured the previously observed distribution of the timescales (Fig. 2b) was preserved. Re-tuning the recurrent connections for the passive task resulted in significantly faster timescales (Supplementary Fig. 6). As expected, the distinct distribution of the input weights projecting to the two inhibitory subpopulations that we observed in Fig. 6a was “flattened” after re-training the DMS RNNs to perform the passive task (Fig. 8e). Repeating the cross-temporal discriminability analysis on the re-trained RNNs showed that the cue stimulus information during the delay period was not maintained as robustly by long *τ* units (Fig. 8f). However, the long *τ* units still carried significantly higher information than the short *τ* units throughout the delay window.

The above results from the experimental data and our model strongly suggest that higher cortical areas might have intrinsically diverse and robust inhibitory signaling. This innate property, in turn, would give rise to long neuronal timescales, and the incoming connections to these areas could undergo plastic changes to support various higher cognitive functions that require integration of information on a slower timescale. Along this line of thought, we hypothesized that the AFC RNNs, which do not have strong inhibitory-to-inhibitory signaling, are not capable of performing WM tasks by simply re-tuning the input weights only. With the recurrent architecture (*W*) fixed, we attempted to re-train the input weights of the 40 AFC RNNs to perform the DMS task, but none of the networks could be trained successfully (yellow line in Fig. 8g). When we repeated the re-training procedure with the *I* → *I* recurrent connections strengthened (see Methods), the performance of the AFC RNNs significantly improved (magenta line in Fig. 8g). On the other hand, the input weights of the DMS RNNs could be successfully tuned to perform the AFC task (Fig. 8h), further confirming the hierarchical organization of these two RNN models.

## Discussion

In this study, we provide a computational model that gives rise to task-specific spontaneous temporal dynamics, reminiscent of the hierarchy of neuronal timescales observed across primate cortical areas [1]. When trained on a WM task, our RNN model was composed of units with long timescales whose distribution was surprisingly similar to the one obtained from the primate dlPFC. In addition, the long-timescale units encoded and maintained WM-related information more robustly than the short-timescale units during the delay period. By analyzing the connectivity structure of the model, we showed that a unique circuit motif that incorporates strong *I* → *I* synapses is an integral component for WM computations and slow baseline temporal properties. Interestingly, *I* → *I* synaptic weights could be manipulated to control both task performance and neuronal timescales tightly. Our work also provides mechanistic insight into how *I* → *I* connectivity supports memory storage and dynamic baseline activity patterns crucial for long neuronal timescales. Lastly, we pro-pose that the microcircuitry we identified is intrinsic to higher-order cortical areas enabling them to perform cognitive tasks that require steady integration of information.

Relating specific baseline spiking activities to the underlying circuit mechanisms has been challenging partly due to the lack of computational models capable of both performing cognitive tasks and capturing temporal dynamics derived from experiments. Although it is possible to use continuous rate RNN models, which have been widely used to uncover neural mechanisms behind cognitive processes [25–29], to study neuronal timescales, our spiking RNN model allowed us to (1) use the same experimental procedures previously used to estimate neuronal timescales, (2) easily interpret and compare our model results with experimental findings, and (3) uncover spiking statistics (spike-count Fano factors) associated with long neuronal timescales.

Our work revealed that strong *I* → *I* connections are critical for long neuronal timescales, and we investigated the functional implication of such connections in WM-related behavior. Despite the fact that excitatory pyramidal cells make up the majority of neurons in cortical areas, inhibitory interneurons have been shown to exert greater influence at the local network level [30, 31]. Further-more, different subtypes of interneurons play functionally distinct roles in cortical computations [9, 14]. In agreement with these observations, recent studies uncovered the importance of disinhibitory gating imposed by VIP interneurons [10, 13, 32, 33]. Through inhibition of SST and PV neurons, VIP interneurons have a unique ability to disinhibit pyramidal cells and create “holes” in a dense “blanket of inhibition” [33]. Surprisingly, optogenetically activating VIP neurons in the PFC of mice trained to perform a WM task significantly enhanced their task performance highlighting that disinhibitory signaling is vital for memory formation and recall [10]. Similar to VIP neurons, SST interneurons have also been shown to disinhibit excitatory cells for fear memory [11, 12]. Intriguingly, the connectivity structures of the RNNs we trained on a WM task using supervised learning also centered around disinhibitory circuitry with strong *I* → *I* synapses (Fig. 6). The strength of the *I* → *I* connections was tightly coupled to the task performance of the RNNs. Thus, our work suggests that microcircuitry specializing in disinhibition could be a common substrate in higher-order cortical areas that require short-term memory maintenance.

Most notably, our results shed light on exactly how robust *I* → *I* connections maintain stable memory storage and long neuronal timescales. By dissecting our WM RNN model, we found that strong mutual inhibition between two oppositely-tuned inhibitory subgroups was necessary for maintaining stimulus-specific information during the delay period (Fig. 6). We also illustrated that our model units that were strongly modulated by *I* → *I* synapses displayed highly dynamic baseline activities leading to both large trial-to-trial Fano factors and long neuronal timescales (Fig. 7). For the neural data, it was not possible to identify neurons tightly regulated by inhibitory synapses, as connectivity patterns are challenging to infer from firing activities alone. However, we showed that long *τ* neurons in the primate PFC also exhibited high trial-to-trial variability. Thus, our findings suggest that spontaneous variability or Fano factors could be a good indicator of the underlying circuit mechanisms: neurons with asynchronously occurring synchronous firing patterns (i.e., high variability) could make up WM-related microcircuits.

Cognitive flexibility is one of the hallmarks of the prefrontal cortex [34, 35]. If higher-order areas are indeed wired with specific and robust *I* → *I* synapses that give rise to stable temporal receptive fields, then what would happen to these connections during learning? Would learning a new task disrupt the existing *I* → *I* connectivity structure, thereby abolishing the previously established timescale distribution? To answer these questions, we analyzed neuronal timescales from monkeys before and after they learned a WM task. The distribution of the timescales was not significantly different between the two conditions, suggesting that learning the WM task did not perturb the intrinsic temporal structure in the dlPFC (Fig. 8). To test if a network is flexible enough to perform other tasks with its timescale dynamics fixed, we re-trained our WM model with its recurrent connectivity frozen to perform a non-WM task (Fig. 8). With re-tuning only the input connections, the WM RNNs could be easily re-trained to perform the non-WM task, implying that these networks are flexible to perform tasks that require not only long timescales but also short timescales. In contrast, RNNs trained on the non-WM task could not be re-trained to perform the WM task. Thus, our work suggests that disinhibitory microcircuits with strong *I* → *I* synapsescould give rise to a flexible module capable of performing a wide range of tasks.

Although our model can capture several experimental findings, a few interesting questions re-main for future studies. For example, our spiking RNN model utilizes connectivity patterns derived from a gradient-descent approach, which is not biologically plausible. It will be important to characterize if more biologically valid learning mechanisms, such as reinforcement learning or Hebbian learning, also generate spiking networks with heterogeneous neuronal timescales. Another unexplored aspect is nonlinear dendritic computations. SST interneurons are known for targeting den-drites of pyramidal cells, and such dendritic inhibition has been associated with gating information [36]. Incorporating dendritic processes into our model could elucidate the computational benefits of dendritic inhibition over perisomatic inhibition during WM. In summary, we have explored a neural circuit mechanism that performs logical computations over time with stable temporal receptive fields.

## Acknowledgements

We are grateful to Ben Tsuda, Yusi Chen, and Jason Fleischer for helpful discussions and feedback on the manuscript. We also thank Jorge Aldana for assistance with computing resources. This work was funded by the National Institute of Mental Health (F30MH115605-01A1 to R.K.). We also gratefully acknowledge the support of NVIDIA Corporation with the donation of the Quadro P6000 GPU used for this research. The funders had no role in study design, data collection and analysis, decision to publish, or preparation of the manuscript.

## Author contributions

R.K. and T.J.S. designed the study and wrote the manuscript. R.K. performed the analyses and simulations.

## Methods

### Continuous rate RNN model

The spiking RNNs used in the main text were generated by first training their counterpart continuous-variable rate RNNs using a gradient descent algorithm. After training, the continuous RNNs were converted to leaky integrate-and-fire (LIF) RNNs using the method that we previously developed [19]. The continuous RNN model contained *N* = 200 recurrently connected units that were governed by

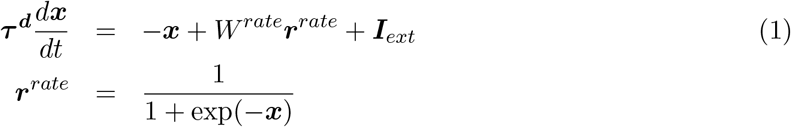

where 20 ms ≤ ***τ***^*d*^ ≤ 125 ms ∈ ℝ^1×*N*^ corresponds to the synaptic decay time constants for the *N* units in the network, ***x*** ∈ ℝ^1×*N*^ is the synaptic current variable, *W* ^*rate*^ ∈ ℝ^*N*×*N*^ is the synaptic connectivity matrix, and ***r***^*rate*^ ∈ ℝ^1×*N*^ refers to the firing rate estimates of the units. A standard logistic sigmoid function was used to estimate a firing rate of a neuron from its synaptic current (*x*).

The external currents (***I***_*ext*_) include task-specific input stimulus signals (see **Training details**) along with a Gaussian white noise variable:

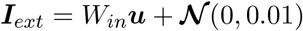

where the time-varying, task-specific stimulus signals 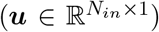 are given to the network via 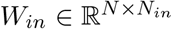, a Gaussian random matrix with zero mean and unit variance. *N*_*in*_ corresponds to the number of input signals associated with a specific task, and 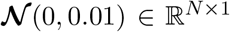 represents a Gaussian random noise with zero mean and variance of 0.01.

A linear readout of the population activity was used to define the output of the rate network:

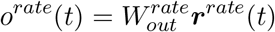

where 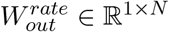 refers to the readout weights.

Eq. (1) is discretized using the first-order Euler approximation method:

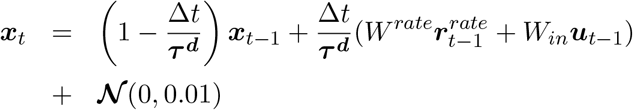

where Δ*t* = 5 ms is the discretization time step size used throughout this study.

### Training details

Adam (adaptive moment estimation), a stochastic gradient descent algorithm, was used to update the synaptic decay variable (*τ*_*s*_), recurrent connections (*W*^*rate*^) and readout weights 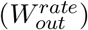. The learning rate was set to 0.01, and the TensorFlow default values were used for the first and second moment decay rates. In addition, Dale’s principle (i.e., separate excitatory and inhibitory populations) was imposed using the method previously proposed [26]. For re-training previously trained RNNs (Fig. 8), only the input weights (*W*_*in*_) were trainable, and the recurrent weights and the readout weights were fixed to their trained values.

Two LIF RNN models were employed in this study by training rate RNNs on two different tasks: delayed match-to-sample (DMS) and 2-alternative forced choice (AFC) tasks.

*DMS RNNs*. For the DMS RNN model, the input matrix (***u*** ∈ ℝ^2×500^) contained two input channels for two sequential stimuli (over 500 time steps with 5 ms step size). The first channel delivered the first stimulus (250 ms in duration) after 1 s (200 time steps) of fixation, while the second channel modeled the second stimulus (250 ms in duration), which began 50 ms after the offset of the first stimulus. The short delay (50 ms) allowed rate RNNs to learn the task efficiently, and the delay duration was increased after training (see below). During each stimulus window, the corresponding input channel was set to either −1 or +1. If the two sequential stimuli had the same sign (−1/−1 or +1/+1), the network was trained to produce an output signal approaching +1 after the offset of the second stimulus. If the stimuli had opposite signs (−1/+1 or +1/−1), then the network produced an output signal approaching −1. The training was stopped when the loss function fell below 7, and the task performance was greater than 95%. After the rate RNNs were successfully trained and converted to LIF networks, a subgroup of LIF RNNs that performed the actual DMS paradigm used in the main text (i.e., delay duration set to 750 ms) with accuracy greater than 95% were identified and analyzed. For Figs. 5, 6 and 7, a group of LIF RNNs that performed the DMS task with accuracy between 60% and 80% was used.

*AFC RNNs*. The input matrix (***u*** ∈ ℝ^1×350^) for the AFC paradigm was set to 0 for the first 200 time steps (i.e., 1 s fixation). A short stimulus (125 ms in duration) of either −1 or +1 was given after the fixation period. After the stimulus offset, the network was trained to produce an output signal approaching −1 for the “−1” stimulus and +1 for the “+1” stimulus. The training termination criteria were the same as those used for the DMS model above.

### Spiking RNN model

For our spiking RNN model, we considered a network of leaky integrate-496 and-fire (LIF) units recurrently connected to one another. These units are governed by:

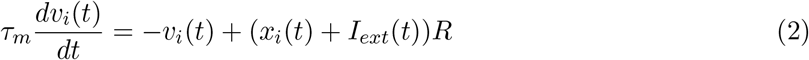

where *τ*_*m*_ is the membrane time constant (10 ms), *v*_*i*_(*t*) is the membrane voltage of unit *i* at time *t*, *x*_*i*_(*t*) is the synaptic input current that unit *i* receives at time *t*, *I*_*ext*_ is the external input current, and *R* is the leak resistance (set to 1). The synaptic input current (*x*) is modeled using a double-exponential synaptic filter applied to the presynaptic spike trains:

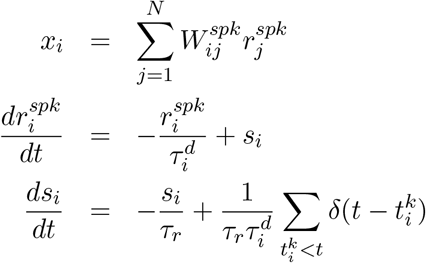

where 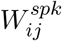 is the recurrent connection strength from unit *j* to unit *i*, *τ*_*r*_ = 2 ms is the synaptic rise time and 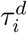 refers to the synaptic decay time for unit *i*. The synaptic decay time constant values and the recurrent connectivity matrix were transferred from the trained rate RNNs (more details described in [19]). The spike train produced by unit *i* is represented as a sum of Dirac *δ* functions, and 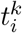 refers to the *k*-th spike emitted by unit *i*.

The external current input (*I*_*ext*_) contained task-specific input values along with a constant background current set near the action potential threshold. The output of our spiking model at time *t* is given by

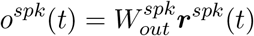

where the readout weights 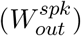 are also transferred from the trained rate RNN model.

Other LIF model parameters included the action potential threshold (−40 mV), the reset potential (−65 mV), the absolute refractory period (2 ms), and the constant bias current (−40 pA). Eq. (2) was discretized using a first-order Euler method with Δ*t* = 0.05 ms.

### Electrophysiological recordings

Extracellular recordings, previously published and described in detail [18, 20, 21], were analyzed to validate our RNN model. The dataset contained spike-train recordings from four rhesus macaque monkeys before and after they learned two DMS tasks. Briefly, for the pre-training condition, the monkeys were rewarded for maintaining fixation on the center of the screen regardless of the visual stimuli shown throughout the trial (Fig. 8a). For the post-training condition, the monkeys were trained on two DMS tasks: spatial and feature DMS tasks. For the spatial task (Fig. 1b), the monkeys were trained to report if two sequential stimuli matched in their spatial locations. For the feature task, they had to distinguish if two sequential stimuli matched in their shapes. The dataset included spike times from single neurons in the dorsal and ventral PFC, but only the units from the dorsal PFC were analyzed for this study.

### Estimation of neuronal timescales

To estimate neuronal timescales, we computed the decay time constant of the spike-count autocorrelation function for each unit during the fixation period [1]. For each unit, we first binned its spike trains during the fixation period over multiple trials using a non-overlapping 50-ms moving window. Since the fixation duration was 1 s for the experimental data and our model, this resulted in a [Number of Trials × 20] spike-count matrix for each unit. For the experimental data, the minimum number of trials required for a neuron to be considered for analysis was 11 trials. The average number of trials from all the neurons from the post-training condition was 86.8 ± 35.1 (mean ± s.d.) trials. For the pre-training condition, the average number of trials was 95.4 ± 44.4. For the RNN model, we generated 50 trials for each unit.

Next, Pearson’s correlation coefficient (*ρ*) was computed between two time bins (i.e., two columns in the spike-count matrix) separated by a lag (∆). The coefficient was calculated for all possible pairs with a maximum lag of 600 ms. The coefficients were averaged for each lag value, and an exponential decay function was fitted across the average coefficient values 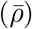 using the Levenberg-Marquardt nonlinear least-squares method:

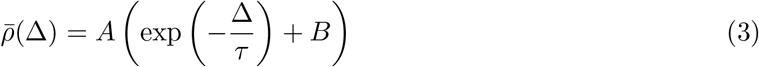

where *A* and *B* are the amplitude and the offset of the fit, respectively. The timescale (*τ*) defines how fast the autocorrelation decays and was used to estimate each neuron’s timescale.

The following inclusion criteria (commonly used in previous experimental studies) were applied to the RNN model and the experimental data: (1) minimum average firing rate of 1 Hz during the fixation period for the experimental data and 2.5 Hz for the RNN model, (2) 0 < *τ* ≤ 500 ms, (3) *A* > 0, and (4) a first decrease in *ρ* earlier than ∆ = 150 ms. In addition, the fitting was started after the first decrease in autocorrelation. For the experimental dataset, 325 dlPFC units from the post-training condition and 434 units from the pre-training condition satisfied the above criteria. For the DMS RNN model, 931 units from 40 good performance RNNs and 604 units from 26 poor performance RNNs met the criteria. For the AFC model, 1138 units from 40 RNNs satisfied the criteria.

### Cross-temporal decoding analysis

The amount of information encoded by each unit was estimated using cross-temporal decoding analysis [8, 22, 23]. For both experimental and model data, a Gaussian kernel (s.d. = 50 ms) was first applied to the spike-trains to obtain the firing rate estimates over time. For each cue stimulus identity, each neuron’s firing rate timecourses were divided into two splits (even vs. odd trials) and averaged across trials within each split. There were 9 cue conditions (i.e., 9 spatial locations) for the spatial DMS task and 8 cue conditions (i.e., 8 shapes) for the feature DMS task. Within each task, all possible pairwise differences in mean firing rates between any two cue conditions for each neuron in each split were computed. Next, Pearson’s correlation coefficient was determined for each pairwise difference condition between the two splits (at each time point across neurons). The correlation coefficients from both tasks (36 pairwise difference conditions for the spatial task and 28 conditions for the feature task) at each time point were averaged after applying the Fisher’s z-transformation resulting in a single measure we refer to as a discriminability or decodability score. The within-time discriminability scores were computed from the correlation coefficients at *t*_1_ = *t*_2_ where *t*_1_ and *t*_2_ refer to the time points used for the two splits.

Nonparametric cluster-based permutation tests were utilized to account for multiple comparisons and to determine significant discriminability (Fig. 3a) and differences in discriminability between short and long *τ* subgroups (Figs. 3 and 8) [37]. To identify significant clusters in the cross-temporal matrices (Fig. 3a and Fig. 8c,f), cue stimulus condition labels were randomly shuffled for 1,000 times within each split to construct the null distribution. A point was considered significant if its value exceeded the 95th percentile of the null distribution, and the largest cluster size (i.e., number of contiguous points that were significant) from the data was compared against the null distribution of the largest cluster size values to correct for multiple comparisons. To determine if within-time decoding timecourses were significantly different between long and short *τ* groups (Fig. 3b and Fig. 8c,f), *τ* group labels were randomly shuffled for 1,000 times within each split and each task. Again, a time point was considered significant if it was greater than the 95th percentile of the null distribution. Similar multiple comparison correction, as described above, was applied.

Cross-temporal decoding matrices and within-time decoding timecourses for the dlPFC data (Figs. 3 and 8) were smoothed for better visualization, but all statistical tests were performed on unsmoothed data.

### Connectivity rewiring method

For Fig. 4e, we characterized which connection type contributed the most to the long neuronal timescales observed in the DMS RNN model by randomly shuffling connections belonging to each type (*I* → *I*, *I* → *E*, *E* → *I*, or *E* → *E*) while preserving the original distribution of the connection types. For the *I* → *I* type, all the outward connections from each inhibitory unit to other inhibitory units were first identified. These connections were then rewired randomly in a manner that preserved their connection identity (i.e., *I* → *I*). This procedure was repeated for the other three synaptic types. For Fig. 5, all the synaptic weights corresponding to each connection type were either decreased or increased by 30% without rewiring.

To quantify the amount of cue-specific information maintained during the delay period in each of the four shuffling conditions (Fig. 4f), we performed the within-time decoding analysis (see above) for all the units in each RNN per shuffling condition. This resulted in 40 within-time decoding timecourses (one for each RNN) for each rewiring condition.

### Cue stimulus selectivity

In order to identify inhibitory units selective for each of the two cue stimuli (−1 or +1), we computed a cue preference index (*θ*) for each unit using:

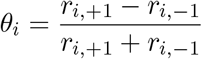

where *r*_*i*,+1_ refers to the average firing rate of unit *i* across positive cue stimulus trials (50 trials) during the cue stimulus window, while *r*_*i*,−1_ indicates the average activity across negative cue stimulus trials (50 trials). Thus, *θ*_*i*_ > 0 indicates that unit *i* prefers the positive cue stimulus over the negative stimulus. Based on this selectivity measure, two subgroups of inhibitory units (one for *θ* > 0 and the other for *θ* < 0) were identified for each DMS RNN.

### Spike-count Fano factors

The relationship between spike-count variability and neuronal timescales was investigated by computing trial-to-trial spike-count Fano factors during the fixation period (Fig. 7). For each unit included in the timescale analysis, the variance of the total number of spikes within the 1-s fixation window across trials was first computed. The Fano factor was then calculated by dividing the variance by the mean spike count. The trials used for computing the Fano factors were identical as those used for estimating the neuronal timescales for both neural and RNN data.

### Reconfiguring pre-trained RNNs

In Fig. 8g,h, the continuous-variable rate RNNs trained to perform the AFC and DMS tasks were used. For Fig. 8g, only the input weights (*W*_*in*_) for the AFC RNNs were re-trained via the same gradient descent algorithm to perform the DMS task. The *I* → *I* connections were either unaltered (yellow in Fig. 8g) or increased by 200% (orange in Fig. 8g). In Fig. 8h, only the input weights for the DMS RNNs were reconfigured to perform the AFC task. The maximum number of training trials was set to 6,000 trials for computational efficiency.

## Code availability

The code for the analyses performed in this work will be made available at https://github.com/rkim35/wmRNN.

## Data availability

The trained RNN models used in the present study will be available as MATLAB-formatted data at https://github.com/rkim35/wmRNN.

## Supplementary Figures

**Supplementary Fig. 1.**
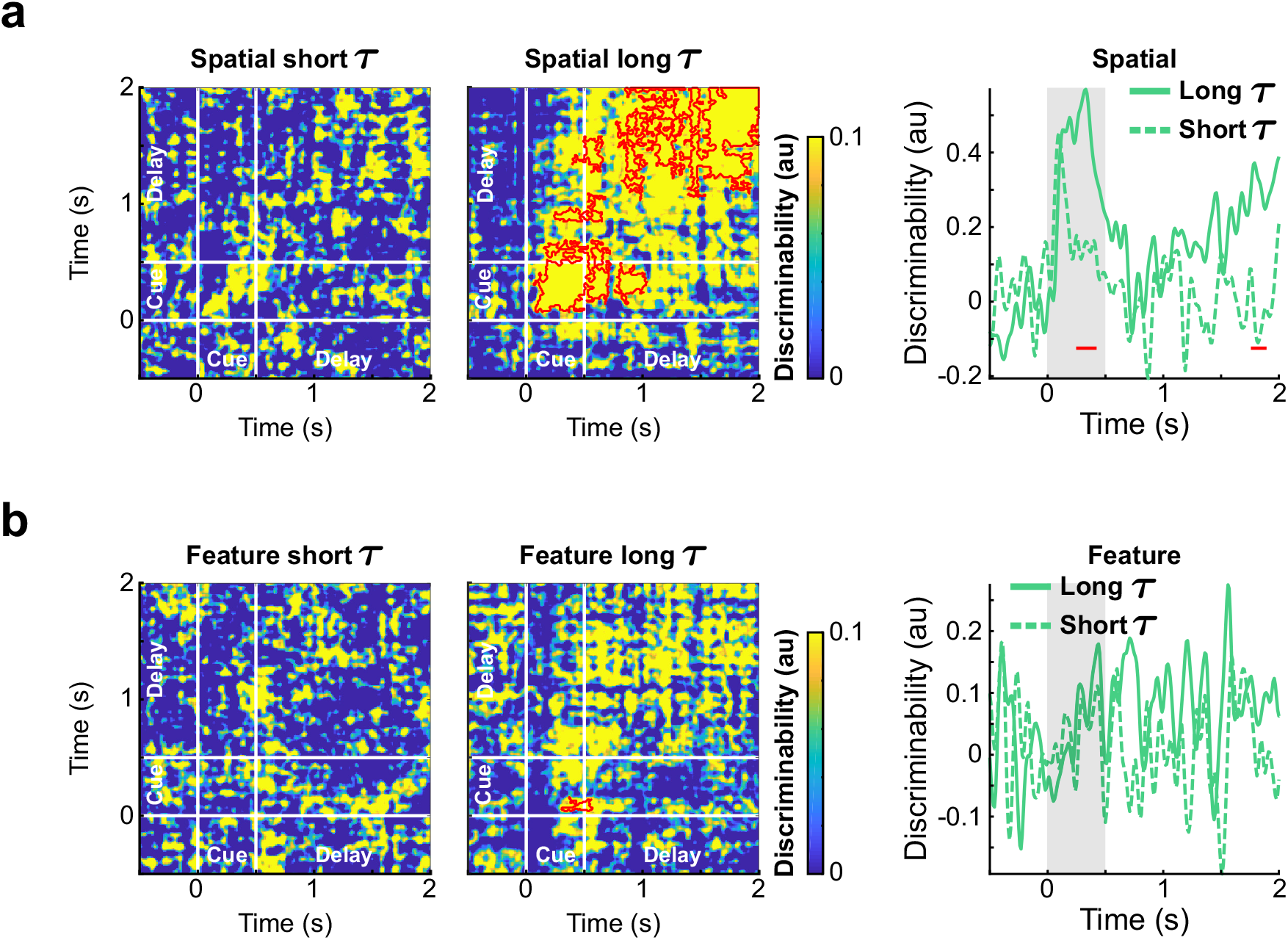
Long *τ* units maintain cue stimulus information during the delay period of the spatial DMS task. **a**, Cross-temporal discriminability matrices and the within-time decoding timecourses from the short and long *τ* groups of the dlPFC data limited to the spatial DMS task. **b**, Cross-temporal discriminability matrices and the within-time decoding timecourses from the short and long *τ* groups of the dlPFC data limited to the feature DMS task. Gray shading, cue stimulus window. Red contours indicate significant decodability (see Methods; cluster-based permutation test, *P* < 0.05). Red lines indicate significant differences in decoding between the short and long *τ* groups (see Methods; cluster-based permutation test, *P* < 0.05).

**Supplementary Fig. 2.**
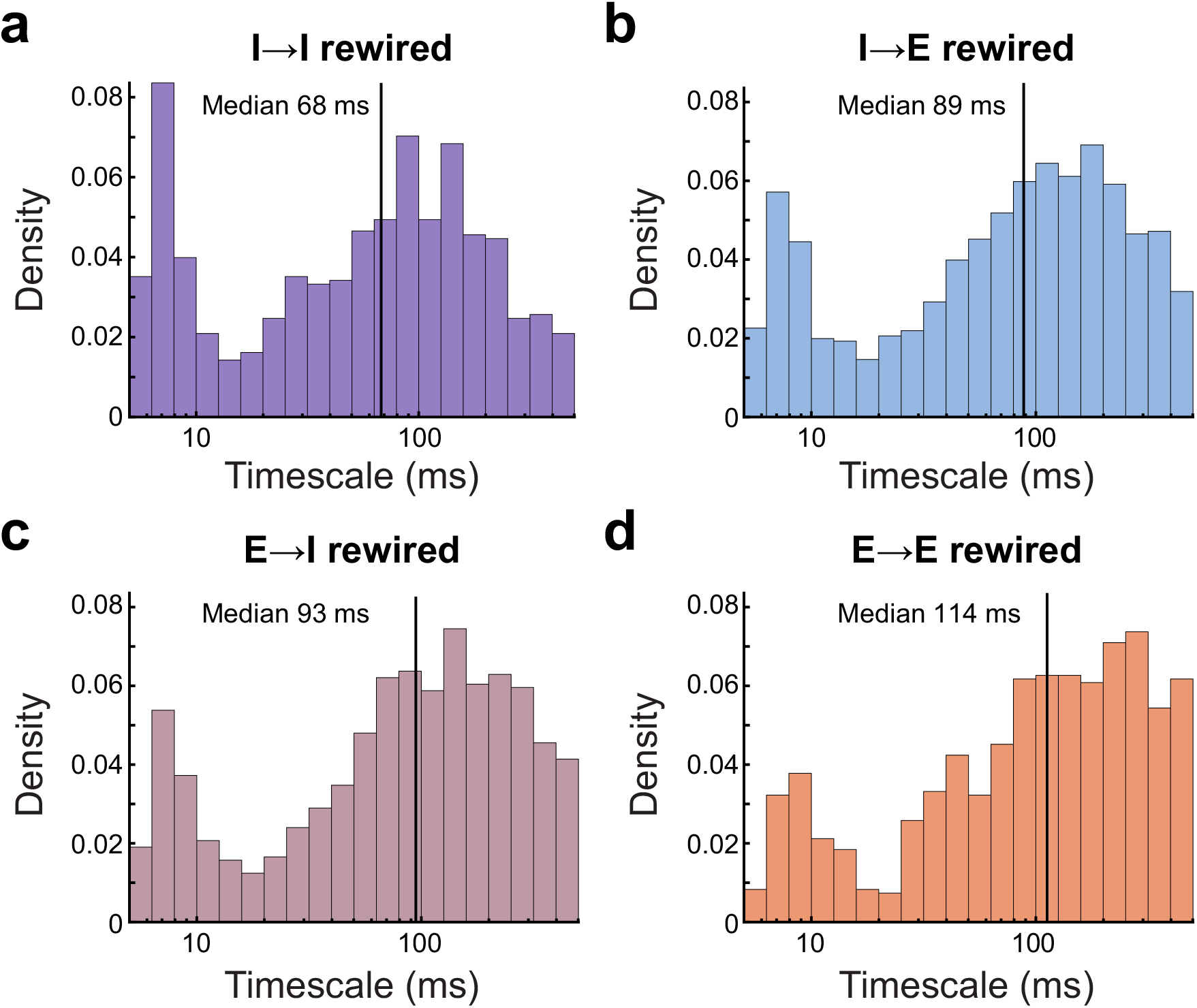
Distribution of the timescales extracted from the DMS RNNs with rewired synaptic connections. **a-c**, Rewiring *I* → *I* (*n* = 824; **a**), *I* → *E* (*n* = 1243; **b**), or *E* → *I* (*n* = 1015; **c**) connections shortened the neuronal timescales. **d**, Shuffling *E → E* (*n* = 891) connections did not alter the distribution significantly. Solid vertical lines represent median log(*τ*).

**Supplementary Fig. 3.**
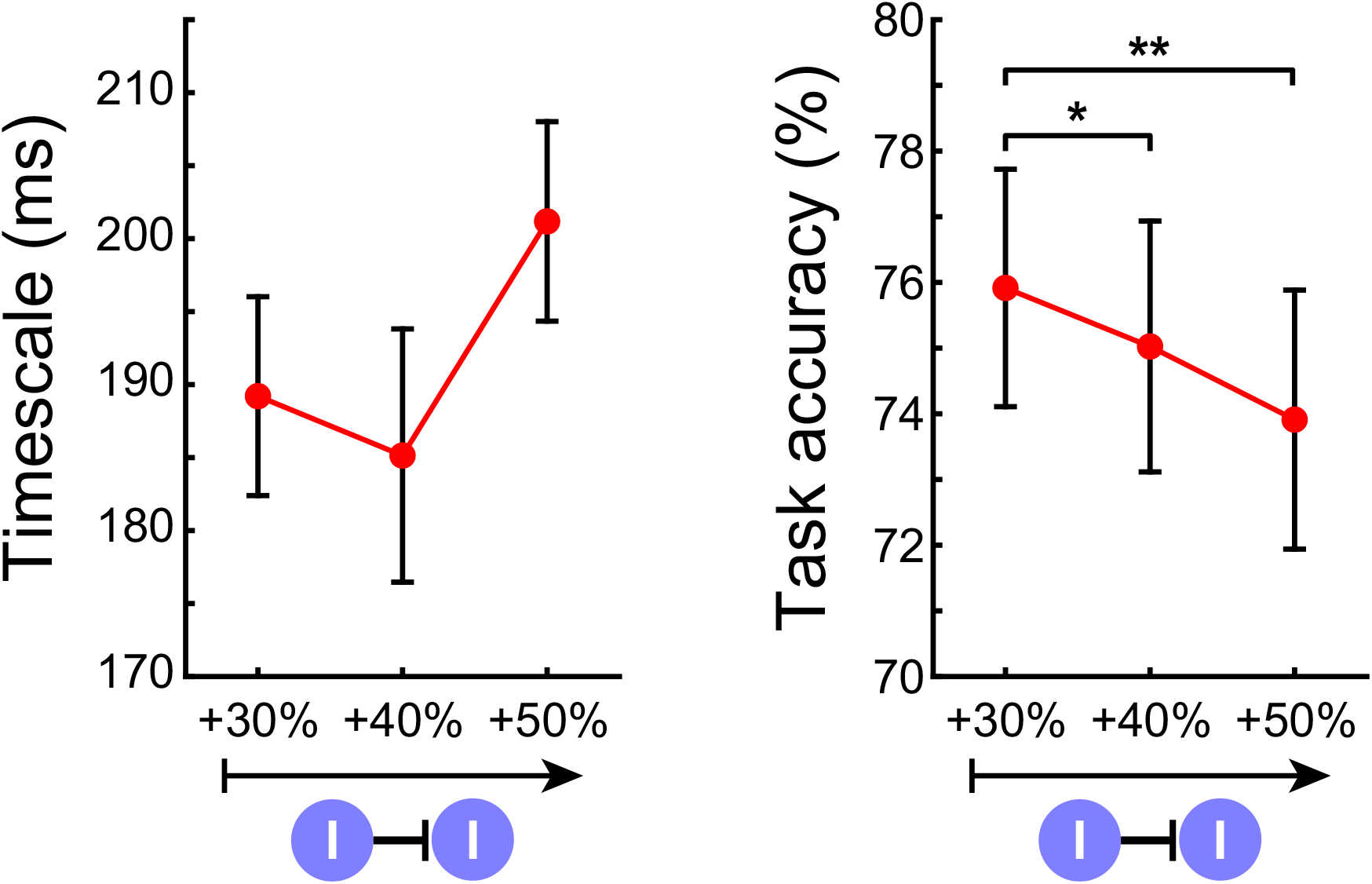
Increasing *I* → *I* connections does not always lead to increased timescales and task performance. Strengthening *I* → *I* connections by more than 30% did not result in significant changes in neuronal timescales (left), but significantly impaired task performance (right). Error bars, ± s.e.m. \**P* < 0.05, \*\**P* < 0.005 by two-sided Wilcoxon rank-sum test.

**Supplementary Fig. 4.**
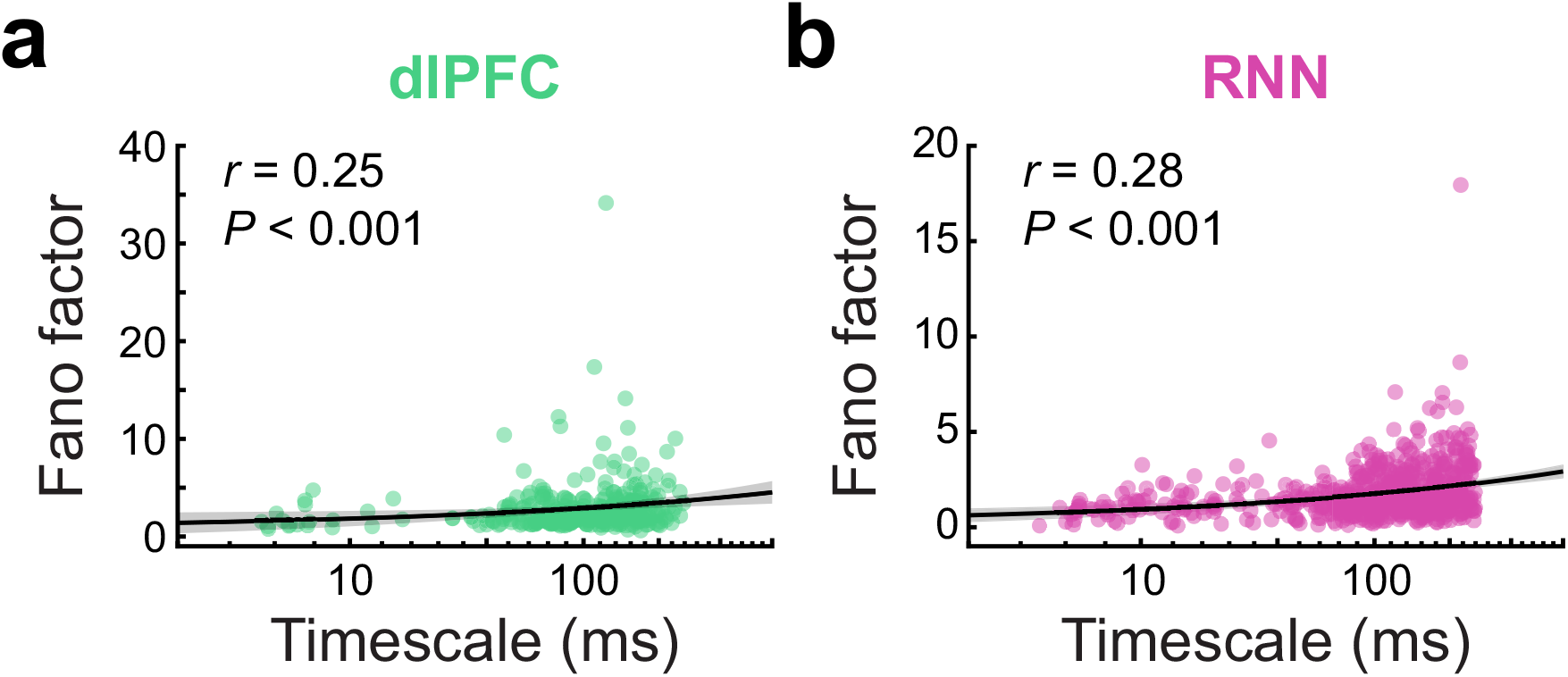
Relationship between trial-to-trial Fano factors and neuronal timescales. **a**,**b**, For both neural (**a**) and RNN data (**b**), trial-to-trial spike-count Fano factors were strongly correlated with neuronal timescales. Spearman rank coefficients are shown. Each dot represents a cell or unit. Black solid lines, linear fits to the log-transformed *τ*; gray shading, 95% confidence interval of the linear fits.

**Supplementary Fig. 5.**
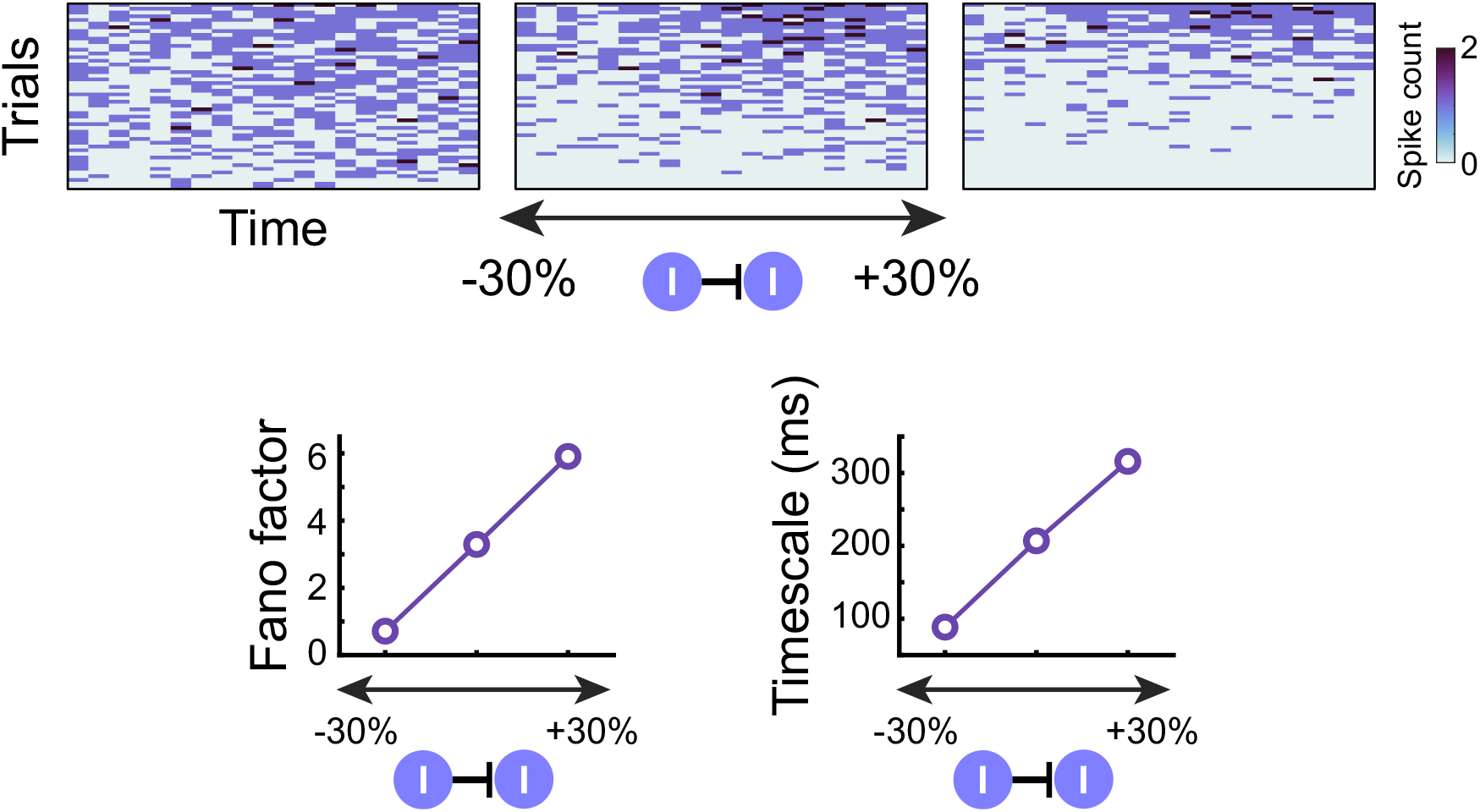
Example unit whose spontaneous firing activity and timescale were strongly modulated by *I* → *I* strength. An inhibitory unit from an example DMS RNN displayed highly dynamic baseline firing activity as *I* → *I* strength increased (top). The unit’s Fano factor and timescale increased linearly with increasing *I* → *I* strength (bottom).

**Supplementary Fig. 6.**
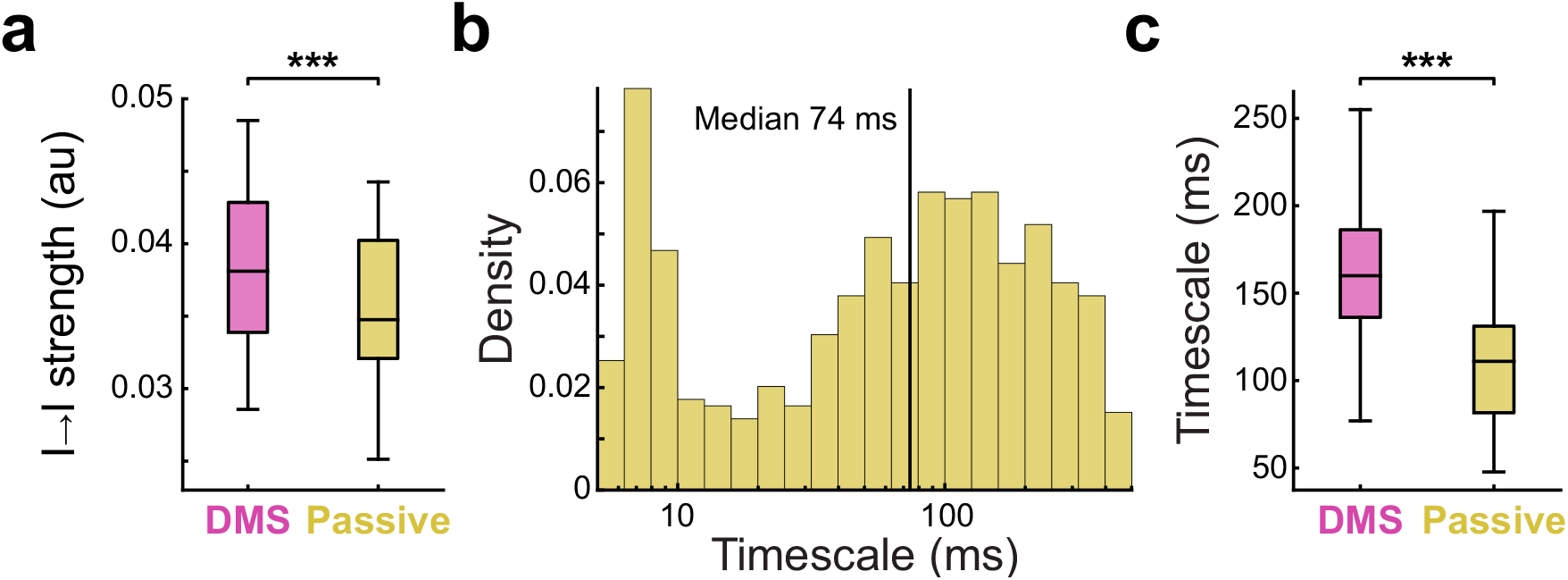
Training DMS RNNs to perform the passive DMS task by retuning recurrent connections (*W*). **a**, Re-tuning the recurrent connections resulted in weakened *I* → *I* connection strength. **b**, Distribution of the timescales (*n* = 598) from the re-trained DMS RNNs. Solid vertical line represents median log(*τ*). **c**, In line with the decreased *I* → *I* strength, the timescales from the re-trained RNNs were significantly shorter than those extracted from the DMS RNNs. Boxplot central lines, median; red circles, mean; bottom and top edges, lower and upper quartiles; whiskers, 1.5*interquartile range; outliers not plotted. \*\*\**P* < 0.0001 by Wilcoxon signed-rank test.

